# Microbial predictors of environmental perturbations in coral reef ecosystems

**DOI:** 10.1101/524173

**Authors:** Bettina Glasl, David G. Bourne, Pedro R. Frade, Torsten Thomas, Britta Schaffelke, Nicole S. Webster

**Author notes:** Address correspondence to Bettina Glasl.

## Abstract

Incorporation of microbial community data into environmental monitoring programs could improve prediction and management of environmental pressures. Coral reefs have experienced dramatic declines due to cumulative impacts of local and global stressors. Here we assess the utility of free-living (i.e. seawater and sediment) and host-associated (i.e. corals, sponges and macroalgae) microbiomes for diagnosing environmental perturbation based on their habitat-specificity, environmental sensitivity and uniformity. We show that the seawater microbiome has the greatest diagnostic value, with environmental parameters explaining 56% of the observed compositional variation and temporal successions being dominated by uniform community assembly patterns. Host-associated microbiomes, in contrast, were five-times less affected by the environment and their community assembly patterns were generally less uniform. Further, seawater microbial community data provided an accurate prediction on the environmental state, highlighting the diagnostic value of microorganisms and illustrating how long-term coral reef monitoring initiatives could be enhanced by incorporating assessments of microbial communities in seawater.

**Importance:** The recent success in disease diagnostics based on the human microbiome has highlighted the utility of this approach for model systems. However, despite improved prediction and management of environmental pressures from the inclusion of microbial community data in monitoring programs, this approach has not previously been applied to coral reef ecosystems. Coral reefs are facing unprecedented pressure on a local and global scale, and sensitive and rapid markers for ecosystem stress are urgently needed to underpin effective management and restoration strategies. In this study, we performed the first assessment of the diagnostic value of multiple free-living and host-associated reef microbiomes to infer the environmental state of coral reef ecosystems. Our results reveal that free-living microbial communities have a higher potential to infer environmental parameters than host-associated microbial communities due to their higher determinacy and environmental sensitivity. We therefore recommend timely integration of microbial sampling into current coral reef monitoring initiatives.

## Introduction

Coral reef ecosystems are rapidly degrading due to local and global pressures (1). Overfishing, pollution, declining water quality, disease and outbreaks of coral predating crown-of-thorns starfish are responsible for localised reef degradation (2) while climate change is impacting reefs on a global scale, including remote reefs with little local anthropogenic pressure (3). For example, elevated sea surface temperatures caused back-to-back coral mass bleaching events in 2016 and 2017, resulting in a significant loss of shallow-water corals on the Great Barrier Reef (GBR) (4). Climate conditions predicted for the end of the century will result in even more frequent and severe coral mass bleaching events with dire projections for the future of coral reefs (5, 6). This global coral reef crisis is driving the development of new management, reef restoration and bioengineering tools to counteract reef loss and ensure the persistence of coral reefs (7, 8). Early prediction of ecosystem stress is critical for an effective implementation of local management and restoration strategies on threatened reef sites.

Microorganisms have considerable potential as a monitoring tool for coral reef ecosystem health (9–11). Microorganisms are fundamental drivers of biogeochemical cycling on coral reefs (12–14), they form intimate associations with the coral reef benthos (15–17), and they contribute significantly to host health and ecosystem homeostasis (18–20). The constant amendment of microbial communities to exploit available resources (21) can trigger differential abundances of specific microorganisms, hence shifts in community composition can provide an early indication of environmental change (22). For example, compositional and functional shifts of coral-associated microbial communities have been described along gradients of anthropogenic impact (23–25) and with changes in water quality (26). However, despite having many of the useful characteristics required of environmental indicators (9, 27), the diagnostic potential of microorganisms for coral reef monitoring is largely conceptual, with only a few studies elaborating on their potential value. For example, the ‘microbialisation score’ measures human impacts on coral reefs based on the ratio of microbial and fish metabolic rates (28). The main limitations to further develop and apply microbial-based monitoring approaches are the lack of temporal and spatial baselines for coral reef microbiomes (9, 29).

Coral reefs comprise a complex network of free-living and host-associated microbial communities with strong benthic-pelagic exchange (13, 30). Therefore, holistic assessments that combine different reef hosts and habitats are required to better understand microbial dynamics and sensitivities to environmental perturbations. The diagnostic value of microbial-based monitoring is likely to vary between distinct habitats of a coral reef ecosystem. For example, microbial communities occurring in seawater may be directly affected by the quality of the ambient reef water or climate conditions, however, the high heterogeneity of seawater due to local hot-spots of available resources (31, 32) may diminish the specificity of these communities. In contrast, microbial communities that dwell in corals live in tight association with the most important frame-builders of reefs (29) and hence may provide crucial information not only on the environmental conditions but also on the effect of the environment on the coral host itself. Sponges, a highly abundant and diverse component of coral reefs (33), are renowned for their enormous filtration capacity (34) and form diverse and intimate associations with microbial communities (35). Hence, sponge microbiomes may provide suitable indicators to monitor water quality. Host-associated biofilms, such as those inhabiting the mucus layer of corals and the surface of macroalgae, provide another potential niche habitat informative for microbial indicators of environmental state. Coral mucus, for example, has been described as a suitable habitat to screen for enterobacteria from sewage contamination due to its ability to trap bacteria (36).

Given the complexity of microbial life on coral reefs we sought to identify the most suitable reef microbiomes for a microbial indicator program to pinpoint environmental state. To do this we quantified the 1) habitat-specificity, 2) determinacy of microbial community successions and 3) sensitivity towards environmental parameters of multiple free-living and host-associated microbiomes. Subsequently, we tested the microbiome’s ability to infer environmental state using indicator value (37) and machine learning approaches (38).

## Results

Samples were collected during a 16-month period (February 2016 - May 2017), at monthly (Magnetic Island - Geoffrey Bay) and seasonal (Orpheus Island – Pioneer Bay – Channel) intervals. The bacterial 16S rRNA genes of 381 samples including seawater, sediment, sponge tissue (*Coscinoderma matthewsi* and *Amphimedon queenslandica*), coral tissue and mucus (*Acropora tenuis* and *Acropora millepora*), and macroalgal surfaces (*Sargassum sp.*) were sequenced (Figure 1). In total 231,316 zero-radius operational taxonomic units (zOTUs) were identified based on 100% sequence similarity (39).

**Figure 1.**
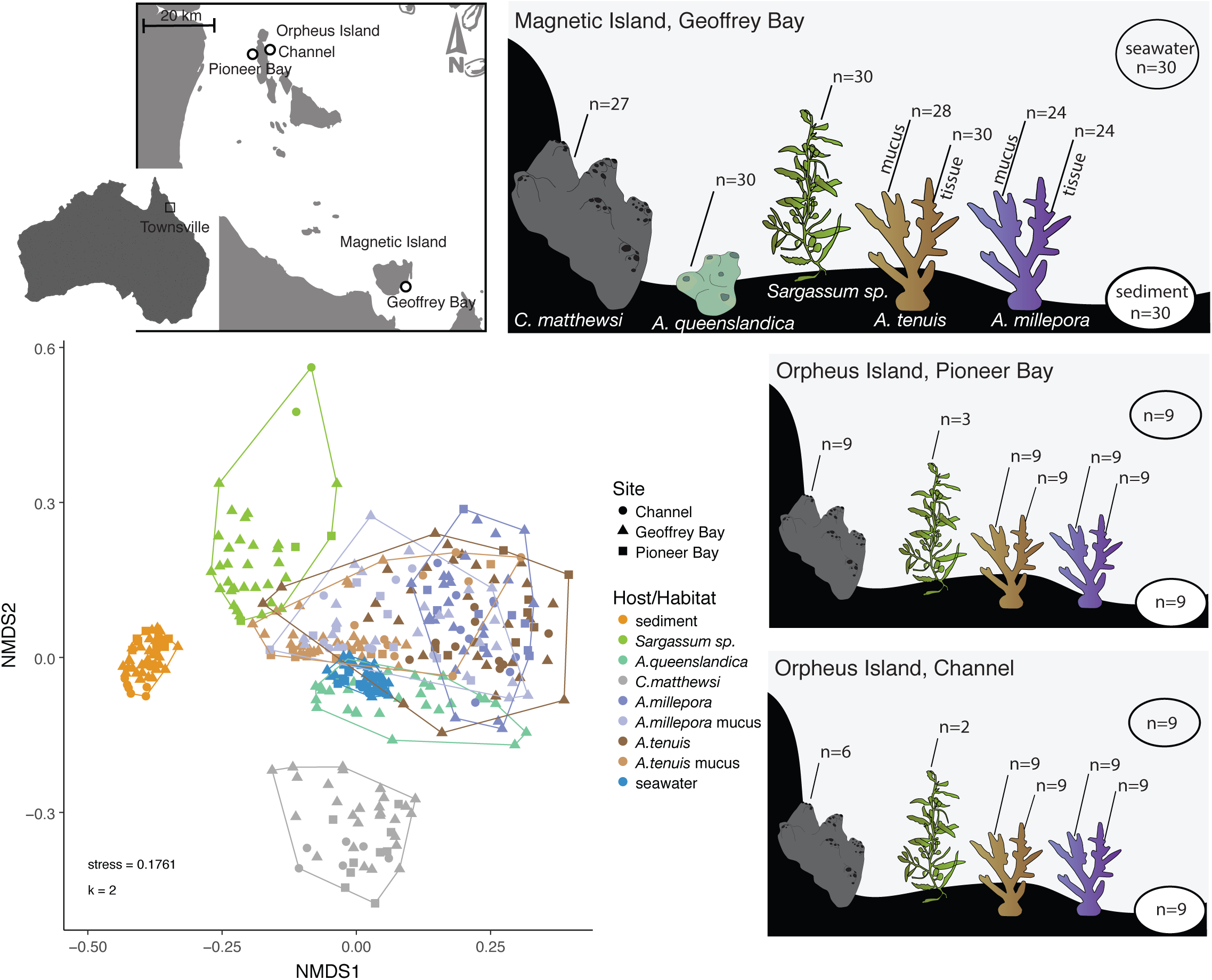
Habitat-specificity of coral reef microbiomes. Seawater, sediment, coral (*Acropora tenuis* and *Acropora millepora*), sponge (*Amphimedon queenslandica* and *Coscinoderma matthewsi*) and macroalgae (*Sargassum sp.*) samples were collected for 16S rRNA gene sequencing at fringing reefs surrounding Magnetic Island (Geoffrey Bay) and Orpheus Island (Pioneer Bay and Channel; Queensland, Australia). Non-metric multidimensional scaling (NMDS) based on Bray-Curtis dissimilarities revealed high habitat-specificity of coral reef microbiomes.

### Coral reef microbiomes are habitat-specific

Habitat-specificity of coral reef microbes was assessed by comparing the similarities of microbial communities associated with seawater (n=48), sediment (n=48), *A. queenslandica* (n=30), *C. matthewsi* (n=42), *A. tenuis* (tissue n=48, mucus n=46*), A. millepora* (tissue n=42, mucus n=42) and *Sargassum sp.* (n=35). Non-metric Multidimensional Scaling based on Bray-Curtis dissimilarities revealed a clear separation of the microbial communities from different reef habitats (Figure 1), and habitat-specificity was further confirmed with Permutational Multivariate Analysis of Variance (PERMANOVA, p = 9.999 × 10^−5^, Table Supplementary Table 1-2). Furthermore, alpha diversities (ANOVA, F_(8/372)_ = 142, p < 2 × 10^−16^) and zOTU richness (ANOVA, F_(8/372)_ = 369, p < 2 × 10^−16^) varied significantly between reef habitats (Supplementary Figure 1 and Supplementary Table 3-5). Sediment harboured by far the most diverse (Shannon Index 7.4 ± 0.2 SD) bacterial community, although microbial diversity was also high in coral surface mucus (Shannon Index 5.1 ± 0.9 SD), macroalgal biofilms (Shannon Index 4.5 ± 1.4 SD), seawater (Shannon Index 4.4 ± 0.2 SD) and in the tissue of the sponge *C. matthewsi* (Shannon Index 4.4 ± 0.3 SD). Microbial diversity was lowest in coral tissue (Shannon Index 3.3 ± 0.8 SD) and in the sponge *A. queenslandica* (Shannon Index 2.7 ± 0.8 SD). These results suggest overall high habitat-specificity of free-living and host-associated microbial communities within coral reef ecosystems.

### Uniform *vs* variable community assembly pattern

The uniformity *versus* variability of microbial community assembly patterns was explored through comparison of compositional similarity (Bray-Curtis index, 0 = dissimilar, 1 = identical) in samples collected monthly at Geoffrey Bay (Magnetic Island). The microbial communities of seawater (n = 30, Wilcoxon Rank-Sum test p =

3.1 × 10^−7^) and sediment (n = 30; Wilcoxon Rank-Sum test p = 3 × 10^−5^) had significantly higher similarities “within” than “among” sampling events (Figure 2a). This uniform response of the free-living microbial communities suggests that deterministic rather than stochastic processes drive their community assembly. For host-associated microbiomes, the overall response pattern varied between species. Microbial communities associated with the sponge *C. matthewsi* (n = 27; Wilcoxon Rank-Sum test, p = 0.0076), the coral *A. tenuis* (mucus n = 28, tissue n = 30; Wilcoxon Rank-Sum test, p = 0.0041 and p = 0.0096, respectively) and the macroalga *Sargassum sp.* (n = 30; Wilcoxon Rank-Sum test, p = 0.00013) followed the same trend as the free-living communities, with significantly higher similarities “within” than “among” sampling events (Figure 2a). In contrast, the microbiome of the sponge *A. queenslandica* (n = 30; Wilcoxon Rank-Sum test, p = 0.23) and the coral *A. millepora* (mucus n = 24, tissue n = 24; Wilcoxon Rank-Sum test, p = 0.15 and p = 0.11 respectively) showed no significant difference in similarities “within” and “among” time points (Figure 2a). Analysis of the compositional similarity of sample replicates within each sampling time point indicated that the seawater microbial communities not only exhibit an overall higher similarity “within” replicates, but the high compositional similarity is conserved across all sampling events (Figure 2b). In contrast, host-associated microbial communities showed a generally lower compositional similarity and higher variation between sample replicates within each sampling time point (Figure 2b).

**Figure 2.**
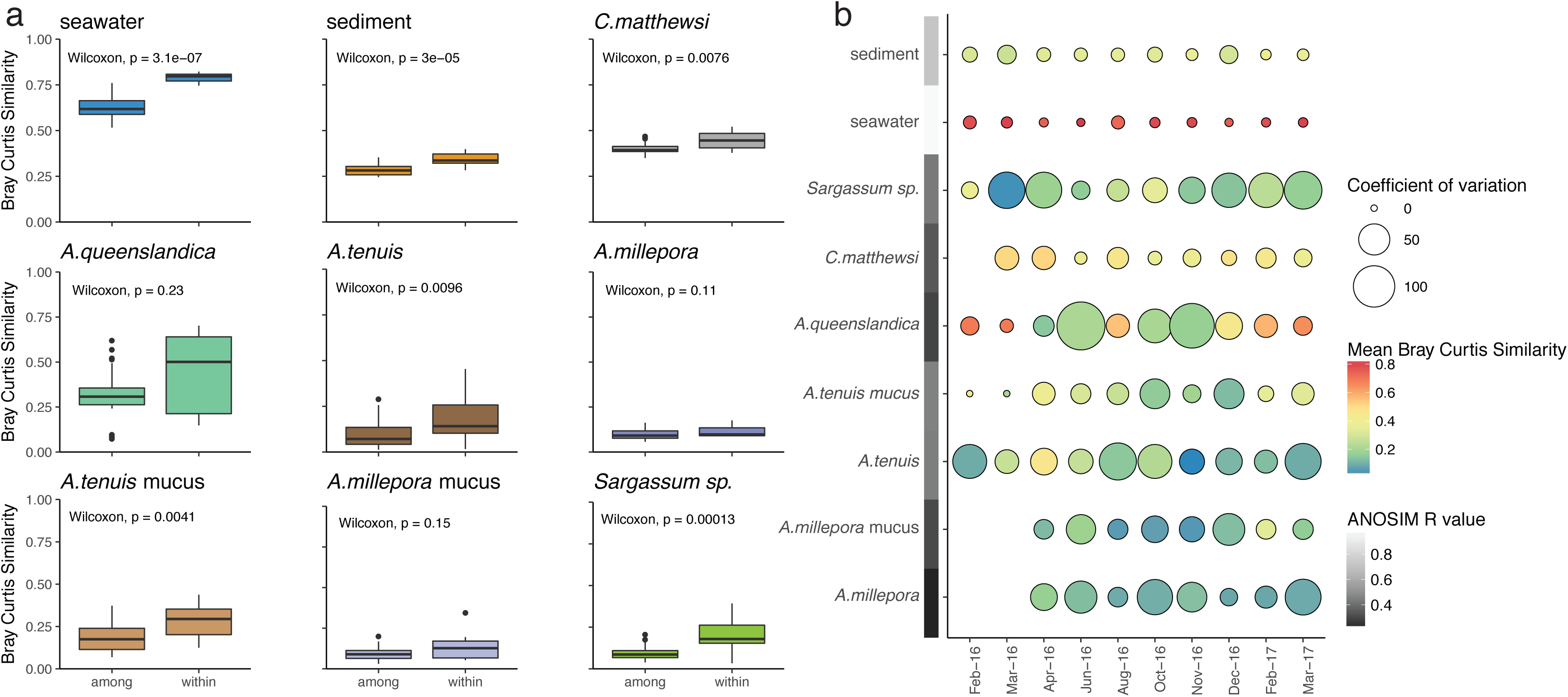
Compositional similarity of coral reef microbiomes over time. a) Variations in the compositional similarity among and within sampling time points of various coral reef microbiomes collected at Geoffrey Bay (Magnetic Island). A higher similarity within time point replicates than among time point replicates suggests a uniform response of the microbial community to temporal variations. Similarities were calculated with Bray-Curtis Similarity Index (0=no similarity, 1=high similarity) and significances tested with Wilcoxon rank-sum test. b) The within sampling time point similarities of replicates (n=3) is indicated in colour and the coefficient of variation (dispersion) is displayed as size. Analysis of Similarity (ANOSIM) was further used as a proxy for the within and among time point variation. R-values of 1 indicate high similarity within sampling time points and high variability among sampling time points, whereas 0 indicates equal similarity within and among sampling time points.

Trends in the temporal community assembly pattern of free-living, host tissue- and biofilm-associated microbial communities were analysed using Analysis of Similarity (ANOSIM) as a proxy to describe similarity patterns (R = 0 indicates equal similarity “within” and “among” time point replicates and R = 1 indicates higher “within” than “among” sampling time point similarities; Figure 2b and Supplementary Figure 2). Overall, free-living microbiomes had R values closer to 1 (seawater R = 0.9919 and sediment R = 0.7322), whereas host tissue-associated microbiomes had R values closer to 0 (*A. queenslandica* R = 0.2927, *C. matthewsi* R = 0.3449, *A. tenuis* tissue R = 0.4547 and *A. millepora* tissue R = 0.2151). Host biofilm-associated microbiomes showed R values of approximately 0.5 (*A. tenuis* mucus R = 0.4613 *A. millepora* mucus R = 0.3090 and *Sargassum sp.* biofilm R = 0.4440). These results suggest that free-living microbiomes (seawater and sediment) exhibit a uniform compositional succession, whereas host-associated microbiomes (coral, sponge and macroalgae) are more stochastic in their temporal community succession. Interestingly, host biofilm-associated microbiomes exhibited a higher uniformity (higher ANOSIM R values) in temporal community succession than tissue-associated microbiomes, most likely reflecting greater environmental influence. The uniform temporal response of free-living microbiomes suggests a high diagnostic value of these microbial communities; hence seawater and sediment microbiomes should provide an accurate prediction of environmental variables.

Microbiomes in seawater (n=48) and sediment (n=48) were further tested for their compositional similarity between all three sampling sites (Geoffrey Bay, Pioneer Bay and Channel). The microbial community composition of sediment samples varied significantly between all three sampling sites (PERMANOVA, p = 9.999 × 10^−5^, 10,000 permutations; Supplementary Figure 3a). The seawater microbiome, in contrast, showed high temporal variability (ANOSIM R = 0.9934, p = 0.001) and low spatial variability (ANOSIM R = 0.2343, p = 0.002; Supplementary Figure 3b). The high spatial variability of sediment microbiomes indicates that habitat characteristics rather than environmental fluctuations are the main drivers structuring community composition.

**Figure 3.**
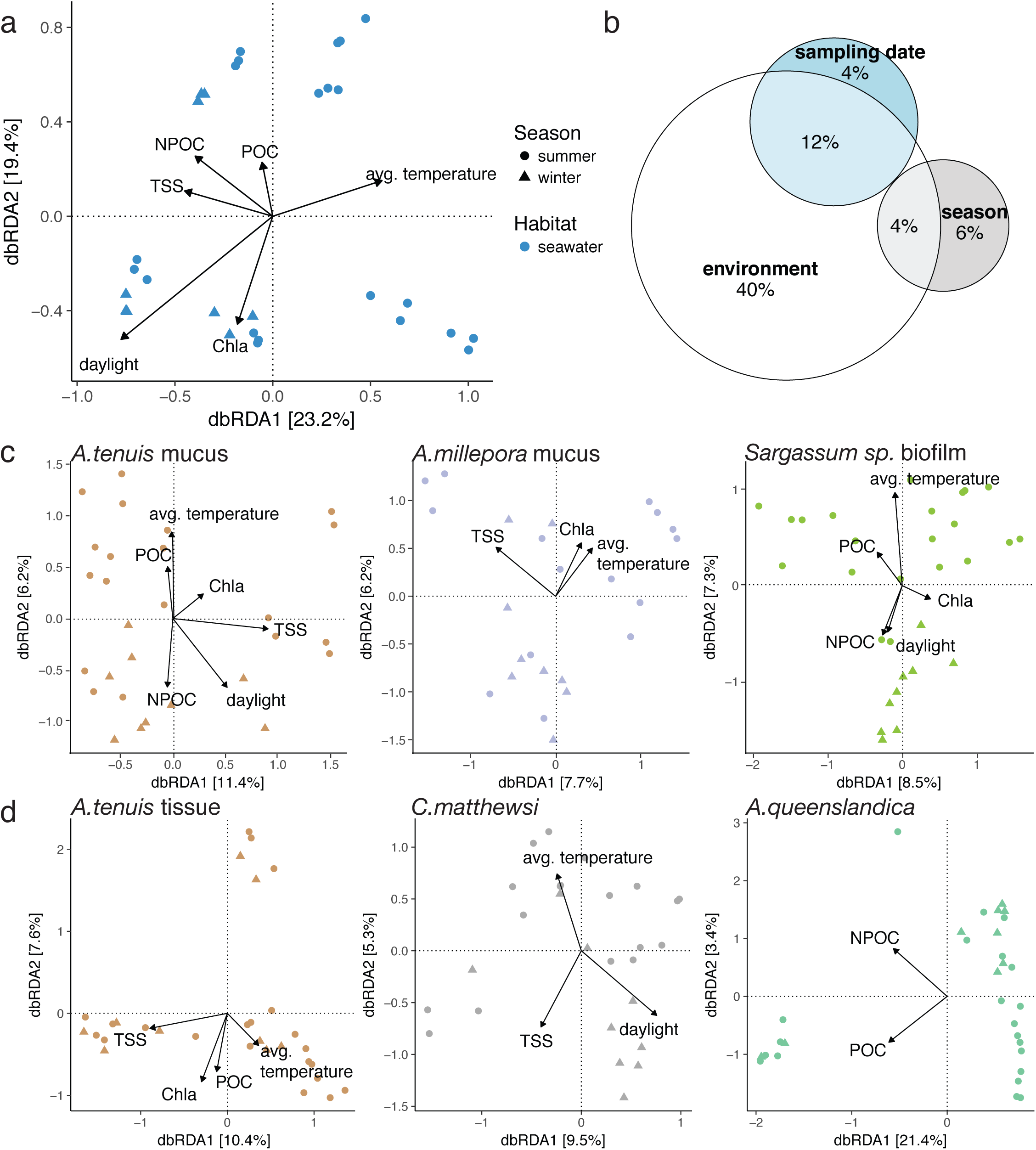
Coral reef microbiome sensitivity to environmental parameters. Bray-Curtis distance-based RDA (dbRDA) was used to evaluate the effect of environmental fluctuations on the microbial community composition of various coral reef habitats/hosts. a) Environmental factors significantly explained 56% of the observed compositional variation in the seawater associated microbial community. b) Variation partitioning shows that environmental parameters rather than season and sampling date explain observed community composition structures in the seawater microbiome. c) Coral mucus and algae biofilm as well as d) coral and sponge tissue microbial communities were significantly influenced by environmental factors; however, environmental parameters only explain on average 11% of the observed community variation.

### Environmental sensitivity

Environmental sensitivity of the different microbiomes was assessed by comparing how much of the compositional variation was explained by sea surface temperature, light and water quality parameters (Supplementary Figures 4 and 5). The compositional variability of the seawater microbiome (n=30) was significantly explained by sampling date, season (summer versus winter) and water quality parameters, such as average seawater temperature, average hours of daylight, total suspended solids (TSS), particulate organic carbon (POC), Chlorophyll *a* (Chl *a*), and non-purgeable organic carbon (NPOC) concentration (PERMANOVA for Bray Curtis distance based Redundancy Analysis (dbRDA); Figure 3a and Supplementary Table 6a-b). In total, these parameters explained 66% of the observed compositional variation in seawater, with 56% being significantly explained by environmental variables (Variation Partitioning Analysis, Figure 3b). Season (summer *versus* winter) and sampling date explained 6% and 4%, respectively (Variation Partitioning Analysis, Figure 3b). In comparison, sampling site significantly explained 24% of the variation in sediment microbial communities (n=48), which overlapped by 12% with the variation explained by sediment characteristics, such as particle size and total organic carbon (TOC) content (PERMANOVA for dbRDA and Variation Partitioning Analysis; Supplementary Table 6b and 7). Water quality parameters and sea surface temperature explained only 3% of the observed variability in the sediment microbiome (Variation Partitioning Analysis).

**Figure 4.**
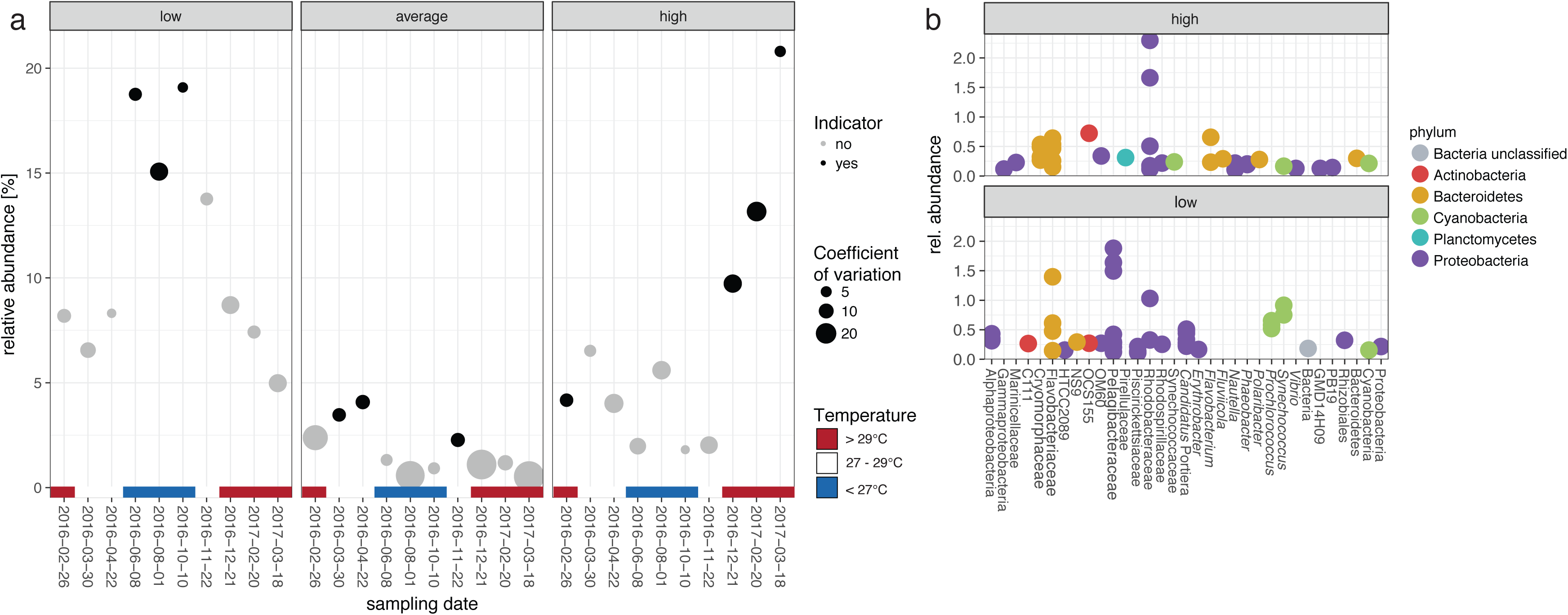
Microbial indicator taxa for seawater temperature fluctuations. Seawater temperatures were z-score standardised and, based on their variation around their mean, classified into low (< −0.5), average (−0.5 – 0.5) and high (> 0.5) temperature groups. Indicator zOTUs were identified with the Indicator Value analysis (IndVal). a) The average relative abundance of the sum of low, average and high temperature indicators is represented for each sampling time point. Significant indicator zOTUs assemblages (p<0.01) for the respective temperature group are indicated in black and size represents the coefficient of variation. Colour gradient further represents the seawater temperature at the given sampling timepoints. b) Relative abundances and taxonomic affiliation of zOTUs identified to be significant (p<0.01) indicators for high and low seawater temperatures. Each dot represents a unique zOTU.

**Figure 5.**
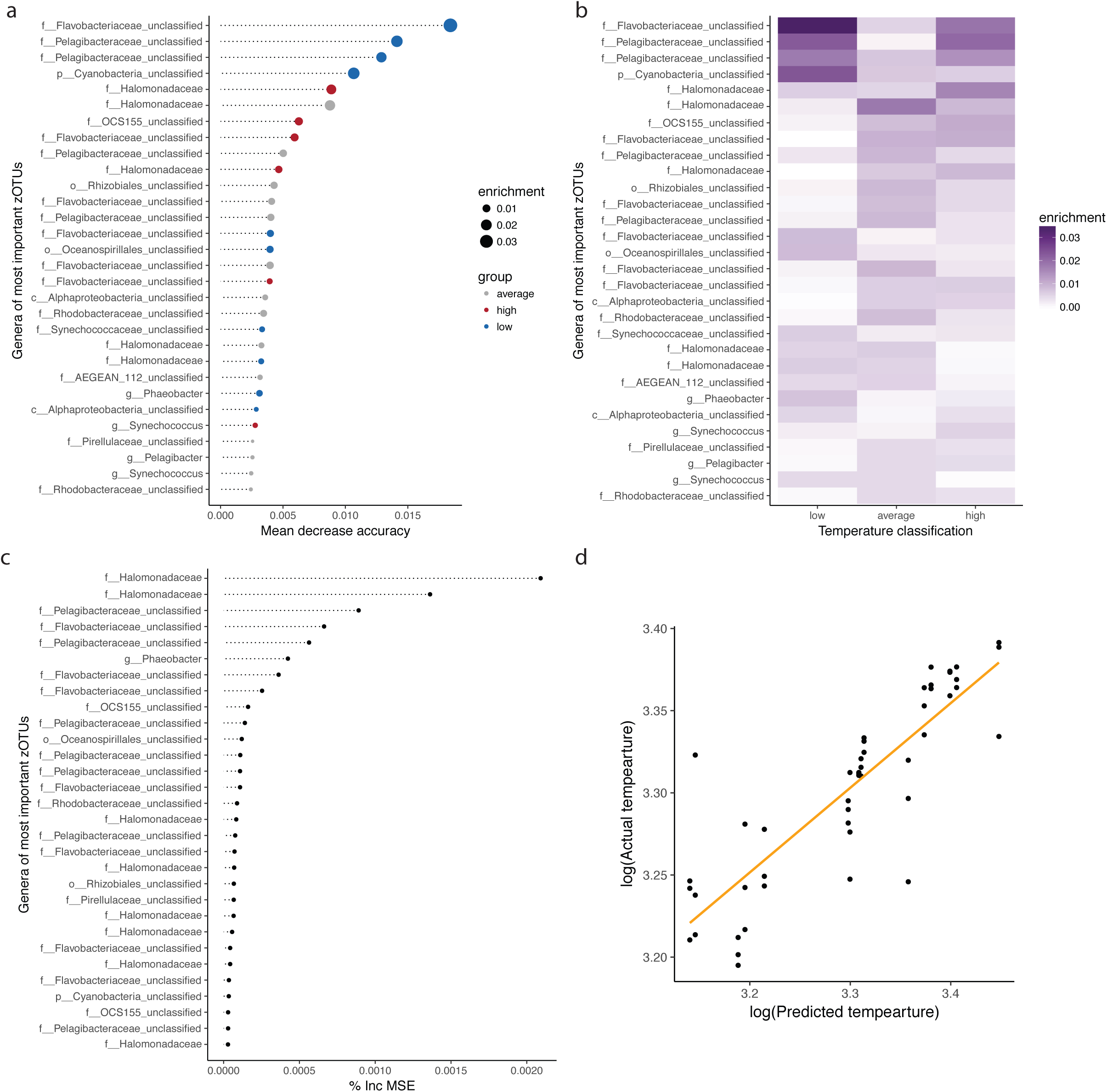
Random Forest machine learning. a) The 30 most important zOTUs reducing the uncertainty in the prediction of seawater temperature classes (low, average, high) based on their mean decrease in accuracy and b) their enrichment in the temperature classes. c) The 30 most important zOTUs reducing the variance (mean squared error (% Inc MSE)) in regression based prediction of seawater temperatures. d) Predicted seawater temperature values *versus* actual seawater temperature values based on Random Forest regression.

Host-associated microbiomes varied substantially in their response to environmental parameters (PERMANOVA for dbRDA and Variation Partitioning Analysis, Figure 3b-c, Supplementary Table 6c-i and 7). On average, 11% of the observed community variations in host-associated microbiomes were explained by the environment, which is five-times less than what we found for the seawater associated microbial community (Supplementary Table 7). This suggests that compositional variations of the seawater microbiome are more likely to reflect environmental changes. Host-associated microbiomes, are comparatively stable to changes in environmental factors.

### Predictability of environmental metadata

Due to the seawater microbiomes uniform temporal pattern and high sensitivity to changing environmental parameters, the ability to infer environmental state based on microbial community data was tested using an Indicator Value analysis (37) and a Random Forest machine learning approach. In total, 110 zOTUs were identified as significant indicators for temperature (Indicator Value p < 0.01). Microbial zOTU assemblages that were indicative of high, low and average seawater temperatures (classification based on their variation around observed annual averages) were present throughout the sampling period. However, higher relative abundances and lower variation (as calculated by coefficient of variation) were evident at certain time points (Figure 4a). Furthermore, we were able to identify microbial indicator taxa for high and low Chl *a*, TSS and POC levels (Supplementary Material Figure 6). Indicators for low and high seawater temperatures were identified in the bacterial phyla Proteobacteria, Bacteroidetes, Cyanobacteria, Actinobacteria and Planctomycetes (Figure 4b). High temperatures were indicated by an increase of zOTUs belonging to the bacterial family *Rhodobacteraceae* and the presence of *Cryomorphaceae*, *Synechococcaeae*, *Vibrio* and *Flavobacterium* (Figure 4b). In contrast, the occurrence of zOTUS belonging to the family *Pelagibacteriaceae* and the genus *Prochlorococcus* were indicative for low seawater temperatures. The phyla Proteobacteria, Bacteroidetes and Cyanobacteria had the greatest number of indicator zOTUs for temperature and other water quality parameters (Supplementary Figure 6). *Flavobacteriaceae*-affiliated zOTUs were significant indicators for temperature, Chl *a*, TSS and POC. *Halomonadaceae* significantly associated with high Chl *a* and TSS and zOTUs belonging to the phylum Verrucomicrobia were significant indicators for high TSS levels.

The diagnostic value of the seawater microbiome (n=48) was further evaluated by applying a Random Forest machine learning classification and regression analysis with 1,213 zOTUs preselected based on a non-zero abundance threshold in at least 10% of the samples (n=48). The seawater microbiome enabled the prediction of seawater temperature classes (low, average, high) with 92% accuracy (Kappa = 88%, Figure 5a-b and Supplementary Figure 7). Highest accuracy (lowest Out of Bag (OOB) estimated error rate) was achieved with m_try_ = 100 zOTUS. Random Forest regression of the seawater microbiome predicted temperature values (R^2^ = 0.67, RMSE = 0.5) (Figure 5c-d and Supplementary Figure 8) with the highest accuracy (lowest OOB estimated error rate) when m_try_ = 400 zOTUs. The effectiveness of zOTUs in reducing uncertainty and variance (also referred to as ‘feature importance’) within the machine learning algorithm was measured by the decrease in mean accuracy for classification and mean-squared error (%incMSE) for regression. The most important zOTUs belong to the bacterial taxa *Flavobacteriaceae*, *Pelagibacteraceae*, Cyanobacteria, *Rhodobacteraceae*, *Synechococcaceae* and *Pirrelulacae*. These results demonstrate that the microbial community associated with coral reef seawater allows for the accurate prediction of fluctuations in sea surface temperature and water quality parameters.

## Discussion

Sensitive and rapidly responding markers of coral ecosystem stress are needed to underpin effective management and restoration strategies. In this study, we used a range of statistical tests and machine learning approaches across multiple free-living and host-associated reef microbiomes to assess their diagnostic value as sensitive indicators of environmental state. Our results show that the microbial community in reef seawater has the highest diagnostic value when compared to other free-living (e.g. sediment) and host-associated microbiomes (e.g. coral, sponge and macroalgae). Our conclusion is based on the microbiome’s 1) habitat-specificity, 2) uniformity of its community assembly, 3) sensitivity towards environmental fluctuations and 4) accuracy to predict environmental parameters. This assessment of the diagnostic capacity of various free-living and host-associated coral reef microbiomes to extrapolate environmental variations provides crucial information for ecosystem management initiatives aimed at incorporating microbial monitoring.

In general, high habitat-specificity was observed across free-living and host-associated microbiomes, confirming previous reports on the compositional variability of microbial communities between coral reef habitats (40), host species (15, 41-43) and even between host compartments (44). High compositional divergence of microbial communities across different reef habitats can be due to the variation of available resources and/or biotic interactions (21). High habitat-specificity contributes to the overall high diversity and complexity across different microbial communities on coral reefs, highlighting the importance of holistic studies that focus on microbial interactions across the benthic-pelagic realm.

Bacterial community structure associated with water and sediment is thought to be primarily governed by deterministic processes (45). Our results are consistent with this, showing uniform community assembly patterns within time point replicates. In contrast, host-associated microbiomes displayed little compositional similarity within a sampling time point, suggesting a non-uniform temporal response. Host-associated microbiomes were also only marginally affected by environmental parameters, indicating that their community assembly pattern are variable between conspecific individuals (45). A higher variability in community assembly can lead to increased community heterogeneity, also referred to as dispersion, which has been described as a common characteristic of host-associated microbiomes (18, 46–48). Furthermore, lower microbial compositional similarities amongst replicates may be driven by increased niche space (e.g. host compartments) (44) and host genotype effects (e.g. host genetics) (42). Collectively, our results show that free-living microbial communities have a higher potential to infer environmental parameters (such as standard measures in environmental monitoring programs) than host-associated microbial communities due to their higher uniformity and environmental sensitivity. Importantly however, previous metaproteomic research on reef sponges has shown that while microbial community composition can appear stable when seawater temperatures increase, disruption to nutritional interdependence and molecular interactions (such as reduced expression of transporters involved in the uptake of sugars, peptides and other substrates) actually occurs prior to detectable changes in community structure (49). Hence, considering the importance of microbes to reef invertebrate health, more sensitive transcriptomic / proteomic approaches may still be warranted for sensitive detection of microbial responses to environmental perturbations.

The diagnostic potential of microbial communities, especially in combination with machine learning approaches, has gained momentum across multiple research fields, including disease identification by characterisation of the human gut-microbiome (50), evaluation of the environment and host genetics on the human microbiome (51), prediction of hydrological functions in riverine ecosystems (52) and assessment of macroecological patterns in soil samples (53). This development of microbial-based diagnostics is largely due to availability of high-throughput sequencing of the 16S rRNA gene and streamlined analytical pipelines that facilitate rapid assessment of microbial community composition (54, 55). In addition to its utility for inferring environmental fluctuations, the seawater microbiome possesses numerous characteristics desirable for environmental monitoring programs: i) non-destructive collection and simple processing methods facilitate large-scale collections alongside existing programs that sample water quality measurements, ii) high fractional contribution of abundant microbes minimises the impacts of sequencing biases (Supplementary Figure 9) and iii) sampling is conducive to future automated, high throughput analyses such as in-line flow cytometry on vessels and real-time DNA/RNA sequencing for community characterisation.

Incorporation of seawater microbial community data into coral reef monitoring approaches should enhance our ability to describe environmental conditions and changes more holistically. For example, temperature fluctuations drive structural variations in seawater microbial communities (56, 57), and elevated seawater temperatures on coral reefs are highly correlated with coral bleaching (1, 58). The inclusion of microbial community data alongside water quality parameters could therefore improve our ability to predict the likelihood of ecosystem stress. For instance, our sample sites, located in the central sector of the GBR, were not affected by the 2016 bleaching that primarily affected the northern sector (59), however they were impacted by the 2017 bleaching event (60). In the months prior to bleaching (late December 2016 till March 2017) we observed two to four times higher relative abundances of high temperature indicator assemblages than when compared to the equivalent period at the beginning of 2016 (Figure 4a), where no bleaching was observed. Interestingly, high temperature indicator assemblages included putative coral pathogens (e.g. *Vibrio*) and opportunistic bacteria (e.g. Rhodobacteraceae, *Verrucomicrobia* and *Flavobacterium*). Coral pathogens, such as *Vibrio corallilyticus* increase their efficiency and motility behaviours with rising seawater temperatures (61–63), and the higher abundance of these microbes may explain the increased prevalence of coral disease post bleaching (64). Hence, microbial monitoring could help inform managers about impending disease outbreaks.

While microbial inventories for reef biofilms and seawater have been established within the Red Sea (57) and Florida coastal areas (65), our study provides the first holistic microbial baseline spanning multiple free-living and host-associated microbiomes for selected GBR sites. Results suggest that there is realistic scope to enhance long-term reef monitoring initiatives by incorporating seawater microbiome observations for assessments of environmental change over space and time, especially for rapid and sensitive identification of early signs of declining ecosystem health. The establishment of microbial observatories (66) and DNA biobanks for long-term biomonitoring (67) will be paramount to successfully inferring ecosystem state and / or perturbations from microbial communities. We therefore recommend timely integration of microbial sampling into current coral reef monitoring initiatives. Further refinement of the sampling and data analysis techniques should focus on selection and validation of additional indicator taxa as well as assessment of ecologically important microbial functions. A further consideration is to explore which monitoring objectives would benefit most from assessments of microbial communities. For example, it is likely that the rapid response time of microbial indicators makes them better suited to early-warning, impact or compliance monitoring programs than to monitoring of slower, long-term changes.

## Materials and Methods

### Sample collection

Samples for microbial community characterization were collected monthly (Magnetic Island) and seasonally (Orpheus Island) from seawater, sediment and multiple host organisms (i.e. corals, sponges and macroalgae), along with environmental metadata, between February 2016 and May 2017 at three Great Barrier Reef sites (Figure 1). Samples were collected under the permit G16/38348.1 issued by the Great Barrier Reef Marine Park Authority.

Samples (n= 3/ sample type/ sampling event) for molecular analysis and additional environmental metadata were collected following the standard operational procedures of the Australian Marine Microbial Biodiversity Initiative (AMMBI; https://data.bioplatforms.com/organization/pages/australian-microbiome/methods). In

brief, seawater for molecular analysis was collected with collapsible sterile bags close to the reef substrate at 2 m depth and pre-filtered (50 µm) to remove large particles and subsequently filtered (2 L) onto 0.2 µm Sterivex-filters (Millepore). The sediment surface layer was sampled with sterile 50 mL tubes at 2 m depth and subsampled immediately into 2 mL cryogenic vials. The sponges *Coscinoderma matthewsi* and *Amphimedon queenslandica* were removed from the substrate (at 7 m and 3 m respectively) with sterile scalpel blades, rinsed with 0.2 µm filter-sterilised seawater and subsampled into 2 mL cryogenic vials. The surface mucus layer of the two acroporid coral species, *Acropora tenuis* and *Acropora millepora*, was sampled with sterile cotton swabs (18). Additionally, coral fragments of each sampled coral were collected at 3 m depth. Coral fragments were rinsed with 0.2 µm filtered-sterilised seawater and placed into 5 mL cryogenic vials. The thallus (including stem, floats and blades) of the macroalgae *Sargassum sp.* was sampled with sterile scalpels at 3 m depth, rinsed with 0.2 µm filtered-sterilised seawater and placed into 2 mL cryogenic vials. All samples were immediately flash frozen in liquid nitrogen after processing and stored at −80°C until DNA extraction.

Additional seawater samples were collected with a diver-operated Niskin bottle close to the reef substrate at 2 m depth at each sampling occasion. Water was subsampled in duplicate for analyses of salinity and concentrations of dissolved organic carbon (DOC), dissolved inorganic carbon (DIC), particulate organic carbon (POC), dissolved inorganic nutrients (DIN), total suspended solids (TSS) and chlorophyll a (Chl a) concentration. Samples were further analysed according to the standard procedures of the Australian Institute of Marine Science (AIMS, Townsville, Australia)(68). Sediment samples were collected with 100 ml glass jars at 2 m depth and characteristics, such as grain size distribution and total organic carbon (TOC) and nitrogen (TON) content, were assessed for each sampling event. Seawater temperatures were obtained from AIMS long-term monitoring temperature records (http://eatlas.org.au/).

### DNA extraction

Prior to extraction, the macroalgal biofilm was separated from the algal tissue by overnight incubation at 200 rpm in 10 mL 1 × PBS at 37°C. Coral fragments were defrosted on ice and the tissue was stripped from the skeleton with an airgun into 1 × PBS solution, homogenised for 1 min at 12.5 rpm with a tissue homogeniser, pelleted (10 min at 16,000 rcf) and snap frozen in liquid nitrogen prior to DNA extraction. DNA from seawater, sediment, sponge and macroalgal biofilms was extracted with the DNeasy PowerSoil kit (Qiagen) and DNA of coral tissue and mucus samples was extracted using the DNeasy PowerBiofilm kit (Qiagen) following the Manufacturer’s instructions. DNA extracts were stored at −80°C until being sent for sequencing.

### 16S rRNA gene sequencing

DNA extracts were sent on dry ice to the Ramaciotti Centre for Genomics (Sydney, Australia) for sequencing. The bacterial 16S rRNA genes were sequenced using the 27F (69) and 519R (70) primer pairs on the Illumina MiSeq platform utilising a duel indexed 2 × 300 bp paired end approach. Further documentation outlining the standard operating procedures for generating and sequencing amplicons is available at https://data.bioplatforms.com/organization/pages/bpa-marine-microbes/methods.

### Sequence analysis

Sequencing data were analysed as single nucleotide variants in a standardized platform alongside other Australian microbial biodiversity initiative samples (39, 71). In brief, forward and reverse reads were merged using FLASH (72). FASTA formatted sequences were extracted from FASTQ files and those < 400 bp in length or containing N’s or homopolymer runs of > 8 bp were removed using MOTHUR (v1.34.1) (73). USEARCH (64 bit v10.0.240) (74) package was used to de-replicate sequences and to order them by abundance. Sequences with < 4 representatives and Chimeras were removed. Quality-filtered sequences were mapped to chimera-free zero-radius operational taxonomic units (zOTUs) and a sample by read abundance table created. zOTUs were taxonomically classified with SILVA v132 (75) database using MOTHUR’s implementation of the Wang classifier (76) and a 60% Bayesian probability cut-off.

Chloroplast and mitochondria derived reads as well as singletons were removed from the dataset. Remaining data were rarefied to 3,600 reads per sample and transformed to relative abundances using the phyloseq package (77) in R (78).

### Habitat and host-specificity

Habitat and host-specificity of a microbiome was assessed by calculating the compositional similarities of all 381 samples with the Bray-Curtis Index and illustrating them in a Non-Metric Multidimensional Scaling (NMDS) plot using the phyloseq package (77). To confirm habitat and host-specificity, Permutational Multivariate Analysis of Variance (PERMANOVA) was applied using the adonis() function of the vegan package (79) with 10,000 permutations.

### Uniform response pattern

The microbiome similarity of replicates for sampling time points *versus* the microbiome similarity among sampling time points was compared by obtaining the Bray-Curtis Similarity for each habitat individually. The variation between the overall within and among time point replicates was tested with a Wilcoxon Rank-sum test in R (78). The dispersion of the Bray-Cutis similarities within a sampling time point was calculated as the coefficient of variation. Analysis of Similarity (ANOSIM; anosim() function of the vegan package (79)) based on Bray-Curtis similarities was used to further evaluate within and among time point similarities in the microbial communities.

### Environmental sensitivities

Environmental metadata were z-score standardized (80) and checked for collinearity using the Pearson correlation coefficient. Collinearity was assumed if correlation was > 0.7 or < −0.7 (81). Collinear variables were considered redundant and removed from the analysis.

zOTU relative abundance, environmental metadata (e.g. average seawater temperature, average hours of daylight, Chl a, POC, NPOC and TSS concentration), season (summer *versus* winter) and sampling date were used for Bray-Curtis distance-based redundancy analysis (db-RDA) using the phyloseq package (77). The significance of each response variable was confirmed with an Analysis of Variance (ANOVA) for the db-RDA (anova.cca() function in the vegan package (79)). Only significant (p-value < 0.05) response variables were kept in the model. The explanatory value (in %) of significant response variables (e.g. environmental parameters, season and sampling date) was assessed with a Variation Partitioning Analysis of the vegan package (79).

### Indicator value analysis

Indicator taxa were identified with the indicator value analysis (indicspecies package (37)) using the following thresholds: 1,000 permutations, minimum specificity (At) and minimum sensitivity (Bt) set to 70% and p-value ≤ 0.01.

### Random forest machine learning

Random forest machine learning was performed with the caret (82) and random forest package (83) in R (78). zOTUs with non-zero abundance values in at least 10% of the samples (n=48) were preselected and z-score standardised prior to model training. Random Forest (with n_tress_ = 10,000) prediction error was measured with out-of-bag (OOB) error. Highest accuracy (lowest OOB estimated error rate) for classification was achieved with m_try_ = 100 zOTUS and for regression with m_try_ = 400 zOTUs. Importance of zOTUs was measured using the decrease in mean accuracy for classification and mean-squared error (%incMSE) for regression.

## Data availability

Sequencing data, metadata and protocols are available at the Bioplatforms Australia data portal under the Australian Microbiome project (www.data.bioplatforms.com). Full usage requires free registration. To search for the sequencing data, navigate to “Processed data”, select “Amplicon is 27f519r_bacteria” and “Environment is Marine”. To search for the Great Barrier Reef sampling sites, add an additional contextual filter, select “Sampling Site” from the dropdown menu and search for “Geoffrey Bay”, “Pionner Bay” and “Channel”.

## Acknowledgements

We thank Michele Skuza, Neale Johnston and the AIMS water quality team for their help with analysing the water quality samples. We also thank Heidi Luter, Katarina Damjanovic and Joe Gioffre for their assistance in the field and Sara Bell for her expertise in the laboratory. We would like to acknowledge the contribution of the Marine Microbes (MM) and Biomes of Australian Soil Environments (BASE) projects, through the Australian Microbiome Initiative in the generation of data used in this publication. The Australian Microbiome Initiative is supported by funding from Bioplatforms Australia through the Australian Government National Collaborative Research Infrastructure Strategy (NCRIS). The study was further funded by the Advance Queensland PhD Scholarship, the Great Barrier Reef Marine Park Authority Management Award and a National Environmental Science Program (NESP) grant awarded to B.G.

## Author contributions

Samples were collected by B.G., D.G.B., P.R.F. and N.S.W. Samples were processed in the laboratory by B.G. and P.R.F. B.G. analysed and prepared the manuscript. All authors reviewed and edited the manuscript.

## References

1. Hughes TP, Barnes ML, Bellwood DR, Cinner JE, Cumming GS, Jackson JBC, Kleypas J, van de Leemput IA, Lough JM, Morrison TH, Palumbi SR, van Nes EH, Scheffer M. 2017. Coral reefs in the Anthropocene. Nature 546:82–90.

2. De’ath G, Fabricius KE, Sweatman H, Puotinen M. 2012. The 27-year decline of coral cover on the Great Barrier Reef and its causes. Proceedings of the National Academy of Sciences of the United States of America 109:17995–17999.

3. Hoegh-Guldberg O, Mumby PJ, Hooten AJ, Steneck RS, Greenfield P, Gomez E, Harvell CD, Sale PF, Edwards AJ, Caldeira K, Knowlton N, Eakin CM, Iglesias-Prieto R, Muthiga N, Bradbury RH, Dubi A, Hatziolos ME. 2007. Coral Reefs Under Rapid Climate Change and Ocean Acidification. Science 318:1737–1742.

4. Hughes TP, Kerry JT, Baird AH, Connolly SR, Dietzel A, Eakin CM, Heron SF, Hoey AS, Hoogenboom MO, Liu G, McWilliam MJ, Pears RJ, Pratchett MS, Skirving WJ, Stella JS, Torda G. 2018. Global warming transforms coral reef assemblages. Nature 556:492–496.

5. van Hooidonk R, Maynard J, Tamelander J, Gove J, Ahmadia G, Raymundo L, Williams G, Heron SF, Planes S. 2016. Local-scale projections of coral reef futures and implications of the Paris Agreement. Sci Rep 6:39666.

6. Hughes TP, Anderson KD, Connolly SR, Heron SF, Kerry JT, Lough JM, Baird AH, Baum JK, Berumen ML, Bridge TC, Claar DC, Eakin CM, Gilmour JP, Graham NAJ, Harrison H, Hobbs JA, Hoey AS, Hoogenboom M, Lowe RJ, McCulloch MT, Pandolfi JM, Pratchett M, Schoepf V, Torda G, Wilson SK. 2018. Spatial and temporal patterns of mass bleaching of corals in the Anthropocene. Science 359:80–83.

7. Anthony K, Bay LK, Costanza R, Firn J, Gunn J, Harrison P, Heyward A, Lundgren P, Mead D, Moore T, Mumby PJ, van Oppen MJH, Robertson J, Runge MC, Suggett DJ, Schaffelke B, Wachenfeld D, Walshe T. 2017. New interventions are needed to save coral reefs. Nat Ecol Evol 1:1420–1422.

8. Damjanovic K, Blackall LL, Webster NS, van Oppen MJH. 2017. The contribution of microbial biotechnology to mitigating coral reef degradation. Microb Biotechnol 10:1236–1243.

9. Glasl B, Webster NS, Bourne DG. 2017. Microbial indicators as a diagnostic tool for assessing water quality and climate stress in coral reef ecosystems. Marine Biology 164.

10. Glasl B, Bourne DG, Frade PR, Webster NS. 2018. Establishing microbial baselines to identify indicators of coral reef health. Microbiology Australia 39:42–46.

11. Roitman S, Joseph Pollock F, Medina M. Coral Microbiomes as Bioindicators of Reef Health, p 1–19 doi:10.1007/13836_2018_29. Springer International Publishing, Cham.

12. Gast GJ, Wiegman S, Wieringa E, Duyl FCv, Bak RPM. 1998. Bacteria in coral reef water types: removal of cells, stimulation of growth and mineralization. Mar Ecol Prog Ser 167:37–45.

13. Bourne D, Webster N. 2013. Coral Reef Bacterial Communities, p 163–187. In Rosenberg E, DeLong E, Lory S, Stackebrandt E, Thompson F (ed), The Prokaryotes doi:10.1007/978-3-642-30123-0_48. Springer Berlin Heidelberg.

14. Sorokin YI. 1973. Trophical role of bacteria in ecosystem of coral reef. Nature 242:415–417.

15. Rohwer F, Seguritan V, Azam F, Knowlton N. 2002. Diversity and distribution of coral-associated bacteria. Mar Ecol Prog Ser 243:1–10.

16. Webster NS, Luter HM, Soo RM, Botte ES, Simister RL, Abdo D, Whalan S. 2012. Same, same but different: symbiotic bacterial associations in GBR sponges. Front Microbiol 3:444.

17. Egan S, Harder T, Burke C, Steinberg P, Kjelleberg S, Thomas T. 2013. The seaweed holobiont: understanding seaweed-bacteria interactions. FEMS Microbiol Rev 37:462–476.

18. Glasl B, Herndl GJ, Frade PR. 2016. The microbiome of coral surface mucus has a key role in mediating holobiont health and survival upon disturbance. ISME J doi:10.1038/ismej.2016.9.

19. Hentschel U, Schmid M, Wagner M, Fieseler L, Gernert C, Hacker J. 2001. Isolation and phylogenetic analysis of bacteria with antimicrobial activities from the Mediterranean sponges Aplysina aerophoba and Aplysina cavernicola. FEMS Microbiol Ecol 35:305–312.

20. Webster NS, Reusch TBH. 2017. Microbial contributions to the persistence of coral reefs. Isme Journal 11:2167–2174.

21. Martiny JB, Jones SE, Lennon JT, Martiny AC. 2015. Microbiomes in light of traits: A phylogenetic perspective. Science 350:aac9323.

22. Garza DR, van Verk MC, Huynen MA, Dutilh BE. 2018. Towards predicting the environmental metabolome from metagenomics with a mechanistic model. Nat Microbiol 3:456–460.

23. Ziegler M, Roik A, Porter A, Zubier K, Mudarris MS, Ormond R, Voolstra CR. 2016. Coral microbial community dynamics in response to anthropogenic impacts near a major city in the central Red Sea. Mar Pollut Bull doi:http://dx.doi.org/10.1016/j.marpolbul.2015.12.045.

24. Kelly LW, Williams GJ, Barott KL, Carlson CA, Dinsdale EA, Edwards RA, Haas AF, Haynes M, Lim YW, McDole T, Nelson CE, Sala E, Sandin SA, Smith JE, Vermeij MJA, Youle M, Rohwer F. 2014. Local genomic adaptation of coral reef-associated microbiomes to gradients of natural variability and anthropogenic stressors. Proceedings of the National Academy of Sciences 111:10227–10232.

25. Dinsdale EA, Pantos O, Smriga S, Edwards RA, Angly F, Wegley L, Hatay M, Hall D, Brown E, Haynes M, Krause L, Sala E, Sandin SA, Thurber RV, Willis BL, Azam F, Knowlton N, Rohwer F. 2008. Microbial ecology of four coral atolls in the northern Line Islands. PLoS ONE 3:e 1584.

26. Angly FE, Heath C, Morgan TC, Tonin H, Rich V, Schaffelke B, Bourne DG, Tyson GW. 2016. Marine microbial communities of the Great Barrier Reef lagoon are influenced by riverine floodwaters and seasonal weather events. PeerJ 4:e1511.

28. Cooper TF, Gilmour JP, Fabricius KE. 2009. Bioindicators of changes in water quality on coral reefs: review and recommendations for monitoring programmes. Coral Reefs 28:589–606.

28. McDole T, Nulton J, Barott KL, Felts B, Hand C, Hatay M, Lee H, Nadon MO, Nosrat B, Salamon P, Bailey B, Sandin SA, Vargas-Angel B, Youle M, Zgliczynski BJ, Brainard RE, Rohwer F. 2012. Assessing coral reefs on a pacific-wide scale using the microbialization score. Plos One 7.

29. Bourne DG, Morrow KM, Webster NS. 2016. Coral Holobionts: Insights into the coral microbiome: Underpinning the health and resilience of reef ecosystems. Annual Reviews of Microbiology 70:317–340.

30. Lesser MP. 2006. Benthic-pelagic coupling on coral reefs: Feeding and growth of Caribbean sponges. Journal of Experimental Marine Biology and Ecology 328:277–288.

31. Azam F. 1998. Microbial control of oceanic carbon flux: the plot thickens. Science 280:694–696.

32. Stocker R. 2012. Marine microbes see a sea of gradients. Science 338:628–633.

33. Diaz MC, Rützler K. 2001. Sponges: an essential component of Caribbean coral reefs. Bull Mar Sci 69:535–546.

34. Reiswig HM. 1971. In situ pumping activities of tropical Demospongiae. Marine Biology 9:38–50.

35. Taylor MW, Radax R, Steger D, Wagner M. 2007. Sponge-associated microorganisms: evolution, ecology, and biotechnological potential. Microbiology and Molecular Biology Reviews 71:295–347.

28. Lipp EK, Griffin DW. 2004. Analysis of coral mucus as an improved medium for detection of enteric microbes and for determining patterns of sewage contamination in reef environments. EcoHealth 1:317–323.

37. De Cáceres M, Legendre P. 2009. Associations between species and groups of sites: indices and statistical inference. Ecology 90:3566–3574.

38. Knights D, Costello EK, Knight R. 2011. Supervised classification of human microbiota. FEMS Microbiol Rev 35:343–59.

39. Brown MV, Kamp Jvd, Ostrowski M, Seymour JR, Ingleton T, Messer LF, Jeffries T, Siboni N, Laverock B, Bibiloni-Isaksson J, Nelson TM, Coman F, Davies CH, Frampton D, Rayner M, Goossen K, Robert S, Holmes B, Abell GCJ, Craw P, Kahlke T, Sow SLS, McAllister K, Windsor J, Skuza M, Crossing R, Patten N, Malthouse P, Ruth PDv, Paulsen I, Fuhrman JA, Richardson A, Koval J, Bissett A, Fitzgerald A, Moltmann T, Bodrossy L. in press. Systematic, continental scale temporal monitoring of marine pelagic microbiota by the Australian Marine Microbial Biodiversity Initiative. Scientific Data SDATA-18-00035B.

40. Tout J, Jeffries TC, Webster NS, Stocker R, Ralph PJ, Seymour JR. 2014. Variability in Microbial Community Composition and Function Between Different Niches Within a Coral Reef. Microb Ecol 67:540–552.

41. Carlos C, Torres TT, Ottoboni LMM. 2013. Bacterial communities and species-specific associations with the mucus of Brazilian coral species. Sci Rep 3.

42. Glasl B, Smith CE, Bourne DG, Webster NS. 2018. Exploring the diversity-stability paradigm using sponge microbial communities. Sci Rep 8:8425.

43. Webster NS, Thomas T. 2016. The Sponge Hologenome. MBio 7.

44. Sweet MJ, Croquer A, Bythell JC. 2011. Bacterial assemblages differ between compartments within the coral holobiont. Coral Reefs 30:39–52.

45. Wang J, Shen J, Wu Y, Tu C, Soininen J, Stegen JC, He J, Liu X, Zhang L, Zhang E. 2013. Phylogenetic beta diversity in bacterial assemblages across ecosystems: deterministic versus stochastic processes. ISME J 7:1310–21.

46. Casey JM, Connolly SR, Ainsworth TD. 2015. Coral transplantation triggers shift in microbiome and promotion of coral disease associated potential pathogens. Sci Rep 5:11903.

47. Zaneveld JR, McMinds R, Thurber RV. 2017. Stress and stability: applying the Anna Karenina principle to animal microbiomes. Nature Microbiology 2.

48. Zaneveld JR, Burkepile DE, Shantz AA, Pritchard CE, McMinds R, Payet JP, RoryWelsh Correa AMS, Lemoine NP, Rosales S, Fuchs C, Maynard JA, Thurber RV. 2016. Overfishing and nutrient pollution interact with temperature to disrupt coral reefs down to microbial scales. Nature Communications 7:11833.

49. Fan L, Liu M, Simister R, Webster NS, Thomas T. 2013. Marine microbial symbiosis heats up: the phylogenetic and functional response of a sponge holobiont to thermal stress. ISME J 7:991–1002.

50. Duvallet C, Gibbons SM, Gurry T, Irizarry RA, Alm EJ. 2017. Meta-analysis of gut microbiome studies identifies disease-specific and shared responses. Nat Commun 8:1784.

51. Rothschild D, Weissbrod O, Barkan E, Kurilshikov A, Korem T, Zeevi D, Costea PI, Godneva A, Kalka IN, Bar N, Shilo S, Lador D, Vila AV, Zmora N, Pevsner-Fischer M, Israeli D, Kosower N, Malka G, Wolf BC, Avnit-Sagi T, Lotan-Pompan M, Weinberger A, Halpern Z, Carmi S, Fu J, Wijmenga C,Zhernakova A, Elinav E, Segal E. 2018. Environment dominates over host genetics in shaping human gut microbiota. Nature 555:210–215.

52. Good SP, URycki DR, Crump BC. 2018. Predicting hydrologic function with aquatic gene fragments. Water Resources Research 54:2424–2435.

53. Ramirez KS, Knight CG, de Hollander M, Brearley FQ, Constantinides B, Cotton A, Creer S, Crowther TW, Davison J, Delgado-Baquerizo M, Dorrepaal E, Elliott DR, Fox G, Griffiths RI, Hale C, Hartman K, Houlden A, Jones DL, Krab EJ, Maestre FT, McGuire KL, Monteux S, Orr CH, van der Putten WH, Roberts IS, Robinson DA, Rocca JD, Rowntree J, Schlaeppi K, Shepherd M, Singh BK, Straathof AL, Bhatnagar JM, Thion C, van der Heijden MGA, de Vries FT. 2018. Detecting macroecological patterns in bacterial communities across independent studies of global soils. Nat Microbiol 3:189–196.

54. Schuster SC. 2008. Next-generation sequencing transforms today’s biology. Nat Methods 5:16–8.

55. Waldor MK, Tyson G, Borenstein E, Ochman H, Moeller A, Finlay BB, Kong HH, Gordon JI, Nelson KE, Dabbagh K, Smith H. 2015. Where next for microbiome research? PLoS Biol 13:e1002050.

56. Sunagawa S, Coelho LP, Chaffron S, Kultima JR, Labadie K, Salazar G, Djahanschiri B, Zeller G, Mende DR, Alberti A, Cornejo-Castillo FM, Costea PI, Cruaud C, d’Ovidio F, Engelen S, Ferrera I, Gasol JM, Guidi L, Hildebrand F, Kokoszka F, Lepoivre C, Lima-Mendez G, Poulain J, Poulos BT, Royo-Llonch M, Sarmento H, Vieira-Silva S, Dimier C, Picheral M, Searson S, Kandels-Lewis S, Tara Oceans c, Bowler C, de Vargas C, Gorsky G, Grimsley N, Hingamp P, Iudicone D, Jaillon O, Not F, Ogata H, Pesant S, Speich S, Stemmann L, Sullivan MB, Weissenbach J, Wincker P, Karsenti E, Raes J,Acinas SG, et al. 2015. Ocean plankton. Structure and function of the global ocean microbiome. Science 348:1261359.

57. Roik A, Rothig T, Roder C, Ziegler M, Kremb SG, Voolstra CR. 2016. Year-Long Monitoring of Physico-Chemical and Biological Variables Provide a Comparative Baseline of Coral Reef Functioning in the Central Red Sea. PLoS One 11:e0163939.

58. Brown EB. 1997. Coral bleaching: causes and consequences. Coral Reefs 16:S129–S138.

59. Hughes TP, Kerry JT, Alvarez-Noriega M, Alvarez-Romero JG, Anderson KD, Baird AH, Babcock RC, Beger M, Bellwood DR, Berkelmans R, Bridge TC, Butler IR, Byrne M, Cantin NE, Comeau S, Connolly SR, Cumming GS, Dalton SJ, Diaz-Pulido G, Eakin CM, Figueira WF, Gilmour JP, Harrison HB, Heron SF, Hoey AS, Hobbs JA, Hoogenboom MO, Kennedy EV, Kuo CY, Lough JM, Lowe RJ, Liu G, McCulloch MT, Malcolm HA, McWilliam MJ, Pandolfi JM, Pears RJ, Pratchett MS, Schoepf V, Simpson T, Skirving WJ, Sommer B, Torda G, Wachenfeld DR, Willis BL, Wilson SK. 2017. Global warming and recurrent mass bleaching of corals. Nature 543:373–377.

60. ARC_Centre_of_Excellence. 2017. Two-thirds of Great Barrier Reef hit by back-to-back mass coral bleaching. Media Release https://www.coralcoe.org.au/media-releases/two-thirds-of-great-barrier-reef-hit-by-back-to-back-mass-coral-bleaching.

61. Garren M, Son K, Raina J-B, Rusconi R, Menolascina F, Shapiro OH, Tout J, Bourne DG, Seymour JR, Stocker R. 2014. A bacterial pathogen uses dimethylsulfoniopropionate as a cue to target heat-stressed corals. The ISME journal 8:999–1007.

62. Garren M, Son K, Tout J, Seymour JR, Stocker R. 2016. Temperature-induced behavioral switches in a bacterial coral pathogen. ISME J 10:1363–72.

63. Tout J, Jeffries TC, Petrou K, Tyson GW, Webster NS, Garren M, Stocker R, Ralph PJ, Seymour JR. 2015. Chemotaxis by natural populations of coral reef bacteria. ISME J 9:1764–1777.

64. Muller EM, Rogers CS, Spitzack AS, van Woesik R. 2008. Bleaching increases likelihood of disease on Acropora palmata (Lamarck) in Hawksnest Bay, St John, US Virgin Islands. Coral Reefs 27:191–195.

65. Campbell AM, Fleisher J, Sinigalliano C, White JR, Lopez JV. 2015. Dynamics of marine bacterial community diversity of the coastal waters of the reefs, inlets, and wastewater outfalls of southeast Florida. Microbiologyopen 4:390–408.

66. Buttigieg PL, Fadeev E, Bienhold C, Hehemann L, Offre P, Boetius A. 2018. Marine microbes in 4D-using time series observation to assess the dynamics of the ocean microbiome and its links to ocean health. Curr Opin Microbiol 43:169–185.

67. Jarman SN, Berry O, Bunce M. 2018. The value of environmental DNA biobanking for long-term biomonitoring. Nat Ecol Evol 2:1192–1193.

68. Devlin MJ, Lourey MJ. 2000. Water quality-field and analytical procedures. Reef L-tMotGB, Australian Institute of Marine Science Townsville.

69. Lane DJ. 1991. 16S/23S rRNA sequencing, p 115–175. In Stackebrandt E, Goodfellow M (ed), Nucleic acid techniques in bacterial systematics. John Wiley and Sons, New York.

70. Turner S, Pryer KM, Miao VP, Palmer JD. 1999. Investigating deep phylogenetic relationships among cyanobacteria and plastids by small subunit rRNA sequence analysis. J Eukaryot Microbiol 46:327–38.

71. Bissett A, Fitzgerald A, Meintjes T, Mele PM, Reith F, Dennis PG, Breed MF, Brown B, Brown MV, Brugger J, Byrne M, Caddy-Retalic S, Carmody B, Coates DJ, Correa C, Ferrari BC, Gupta VVSR, Hamonts K, Haslem A, Hugenholtz P, Karan M, Koval J, Lowe AJ, Macdonald S, McGrath L, Martin D, Morgan M, North KI, Paungfoo-Lonhienne C, Pendall E, Phillips L, Pirzl R, Powell JR, Ragan MA, Schmidt S, Seymour N, Snape I, Stephen JR, Stevens M, Tinning M, Williams K, Yeoh YK, Zammit CM, Young A. 2016. Introducing BASE: the Biomes of Australian Soil Environments soil microbial diversity database. Gigascience 5.

72. Magoc T, Salzberg SL. 2011. FLASH: fast length adjustment of short reads to improve genome assemblies. Bioinformatics 27:2957–63.

73. Schloss PD, Westcott SL, Ryabin T, Hall JR, Hartmann M, Hollister EB, Lesniewski RA, Oakley BB, Parks DH, Robinson CJ, Sahl JW, Stres B, Thallinger GG, Van Horn DJ, Weber CF. 2009. Introducing mothur: open-source, platform-independent, community-supported software for describing and comparing microbial communities. Appl Environ Microbiol 75:7537–41.

74. Edgar RC. 2010. Search and clustering orders of magnitude faster than BLAST. Bioinformatics 26:2460–1.

75. Yilmaz P, Parfrey LW, Yarza P, Gerken J, Pruesse E, Quast C, Schweer T, Peplies J, Ludwig W, Glockner FO. 2014. The SILVA and “All-species Living Tree Project (LTP)” taxonomic frameworks. Nucleic Acids Res 42:D643–8.

76. Wang Q, Garrity GM, Tiedje JM, Cole JR. 2007. Naïve bayesian classifier for rapid assignment of rRNA sequences into the new bacterial taxonomy. Appl Environ Microbiol 73:5261–5267.

77. McMurdie PJ, Holmes S. 2013. phyloseq: an R package for reproducible interactive analysis and graphics of microbiome census data. PLoS One 8:e61217.

78. R Development Core Team. 2008. R: A language and environment for statistical computing. R Foundation for Statistical Computing.

79. Oksanen J, Blanchet FG, Kindt R, Legendre P, Minchin PR, O’Hara RB, Simpson GL, Solymos P, Stevens MHH, Wagner H. 2013. vegan: Community Ecology Package. R package version 20–9.

80. Clark? Carter D (ed). 2014. z Scores. Wiley StatsRef.

81. Dormann CF, Elith J, Bacher S, Buchmann C, Carl G, Carre G, Marquez JRG, Gruber B, Lafourcade B, Leitao PJ, Munkemuller T, McClean C, Osborne PE, Reineking B, Schroder B, Skidmore AK, Zurell D, Lautenbach S. 2013. Collinearity: a review of methods to deal with it and a simulation study evaluating their performance. Ecography 36:27–46.

82. Kuhn M. 2008. Caret package. Journal of Statistical Software 28.

83. Liaw A, Wiener M. 2002. Classification and Regression by randomForest. R News 2:18–22.

